# sigLASSO: optimizing cancer mutation signatures jointly with sampling likelihood

**DOI:** 10.1101/366740

**Authors:** Shantao Li, Forrest W. Crawford, Mark B. Gerstein

## Abstract

Multiple mutational processes drive carcinogenesis, leaving characteristic signatures on tumor genomes. Determining the active signatures from the full repertoire of potential ones can help elucidate mechanisms underlying cancer initiation and development. This task involves decomposing the counts of cancer mutations, tabulated according to their trinucleotide context, into a linear combination of known mutational signatures. We formulate it as an optimization problem and develop sigLASSO, a software tool, to carry it out efficiently. (An R package implementation is available at github.com/gersteinlab/siglasso). sigLASSO features four key aspects: (1) It jointly optimizes the likelihood of sampling and signature fitting, by explicitly adding multinomial sampling into the overall objective function. This is particularly important when mutation counts are low and sampling variance is high, such as in exome sequencing. (2) sigLASSO uses L1 regularization to parsimoniously assign signatures to mutation profiles, leading to sparse and more biologically interpretable solutions resembling previously well-characterized results. (3) sigLASSO fine-tunes model complexity, informed by the scale of the data and biological-knowledge based priors. In particular, instead of hard thresholding and choosing a priori a discrete subset of active signatures, sigLASSO allows continuous priors, which can be effectively learned from auxiliary information. (4) Because of this, sigLASSO can assess model uncertainty and abstain from making certain assignments in low-confidence contexts. Finally, to evaluate sigLASSO signature assignments in comparison to other approaches, we develop a set of reasonable expectations (e.g. sparsity, the ability to abstain, and robustness to noise) that we apply consistently in a variety of contexts.

## 1 Introduction

Mutagenesis is a fundamental process underlying cancer development. Examples of mutational mechanisms include spontaneous deamination of cytosines, the formation of pyrimidine dimers by ultraviolet (UV) light, and the crosslinking of guanines by alkylating agents. Multiple endogenous and exogenous mutational processes drive cancer mutagenesis and leave distinct fingerprints (1). Notably, these processes have characteristic mutational nucleotide context biases. (2; 3; 4; 5; 6) Sequencing cancer samples at presentation revealed all mutations accumulated over lifetime; these include somatic alterations generated by multiple mutational processes both before cancer initiation and during cancer development. In a generative model, multiple latent mutational processes generate mutations over time, drawing from their corresponding nucleotide context distributions (“mutation signature”). (4; 5) Here, a mutation signature is a multinomial probability distribution of mutations of a set of nucleotide contexts. In cancer samples, mutations from various mutational processes are mixed and observable by sequencing.

By applying unsupervised methods such as non-negative matrix factorization (NMF) and clustering to large-scale cancer studies, researchers have decomposed the mutation mixture and identified at least 30 distinct mutational signatures. (2; 7) Many signatures have been linked with mutational processes with known etiologies, such as aging, smoking, or ApoBEC activity. Investigating the fundamental processes underlying mutagenesis could help elucidate the initiation and development of cancer.

A major task in cancer research is to leverage signature studies on large-scale cancer cohorts and efficiently select and attribute active signatures to new cancer samples. A popular previously published method, deconstructSigs, (8) decomposes the mutation profile into a signature mixture using binary search to iteratively test coefficients one-by-one and then hard pruning signatures with low estimated contribution to achieve sparsity. Other approaches use linear programming (9) or iterate all combinations by brute force.(10) None of these approaches explicitly formulates sampling uncertainty into the model or uses efficient regression techniques. Moreover, no off-the-shelf implementation of these methods, besides deconstructSigs, is available.

Although we do not fully know the latent mutational processes in cancer samples, we can make reasonable and logical assumptions that facilitate our method design. Here, we aimed to design a computational framework, sigLASSO, that could meet these criteria. First, we assumed that the set of estimated mutational mechanisms should be small, as *de novo* studies indicate that not all signatures can be active in a single sample or even a given cancer type. In most cancer samples, only a few signatures are identified in original *de novo* discovery studies. An ideal tool should generate solutions that follow this sparse distribution. Moreover, the number of detectable mutation signatures is limited by the amount of data support. Too many signatures leads to overfitting and unstable solutions. We aimed for a sparser solution as it explains observations in a simpler fashion. Second, the estimated mutational mechanisms should be biologically interpretable and reflect some cancer type specificity. For example, we should not observe UV-associated signatures in tissues that are not exposed to UV. Likewise, we only expect to observe activation-induced cytidine deaminase mutational processes, which are biologically involved in antibody diversification, in B-cell lymphomas. Finally, we felt the solution should be robust and the data should control the model complexity.

In particular, reliably recovering the signature composition is challenging when the mutation number is low. (8) Low mutation count results in high sampling variance, leading to an unreliable estimation of the mutation context probability distribution, which is the target for signature fitting. A desirable signature identification tool should model the sampling process and take sampling variance into consideration.

In this work, we formulated the task as a joint optimization problem with L1 regularization. First, by jointly fitting signatures and the parameters of a multinomial sampling process, sigLASSO takes into account the sampling uncertainty. Cooperatively fitting a linear mixture and maximizing the sampling likelihood enables knowledge transfer and improves performance. Specifically, signature fitting imposes constraints on the previously unconstrained multinomial sampling probability distribution. Conversely, a better estimation of the multinomial sampling probability improves signature fitting. This property is especially critical in high sampling variance settings, for example, when we only observe low mutation counts in whole exome sequencing (WES). Existing methods use continuous relaxation, which makes the model invariant to different mutation counts. Second, sigLASSO penalizes the model complexity and achieves variable selection by regularization. Regularization is essential for this fitting problem with a large amount signatures to achieve proper variable selection and avoid overfitting. Using regularization to promote sparsity and prevent overfitting is also a standard practice in *de novo* signature discovery. A few recent examples are rule-based constrains (SigProfiler), a Bayesian variant of NMF that penalizes on model complexity (SignatureAnalyzer, (11)) and L1 based regularization (SparseSignatures, (12)). The most straightforward way to do this would be to use the L0 norm (cardinality of active signatures), but this approach cannot be effectively optimized. Conversely, using the L2 norm flattened out at small values leads to many tiny, non-zero coefficients, which do not resemble the sparse signature distribution in the original *de novo* studies and are hard to interpret biologically. sigLASSO uses the L1 norm, which promotes sparsity. The L1 norm is convex, and thus allows efficient optimization. (13; 14) Additionally, this approach is able to harmoniously integrate prior biological knowledge into the solution by fine-tuning penalties on the coefficients. Compared with the approach of subsetting signatures before fitting, our soft thresholding method is more flexible to noise and unidentified signatures. Finally, sigLASSO is aware of data complexity such as mutational number and patterns in the observation. It is able to abstain (decline to assign mutational processes under high uncertainty) and defer to the human researcher to decide. Our method is automatically parameterized empirically on performance, allowing data complexity to inform model complexity. In this way, our approach also promotes result reproducibility and fair comparison of datasets.

In sum, sigLASSO exploits constraints in signature identification and provides a robust framework for scientists to achieve biologically sound solutions. sigLASSO also can empower researchers to use and integrate their biological knowledge and expertise into the model. Unveiling the underlying mutational processes in cancer samples will enable us to recognize and quantify new mutagens, understand mutagenesis and DNA repair processes, and develop new therapeutic strategies for cancer. (9; 15; 16; 17; 18)

## 2 MATERIALS AND METHODS

### The signature identification problem

Mutational processes leave mutations in the genome within distinct nucleotide contexts. We denoted the total number of contexts as *n*. Typically, we considered the mutant nucleotide context and looked one nucleotide ahead and behind each mutation, dividing the mutations into *n* = 96 trinucleotide contexts. Each mutational process carries a unique signature, which is represented by a multinomial probability distribution over mutations of trinucleotide context (Fig 1a).

**Figure 1:**
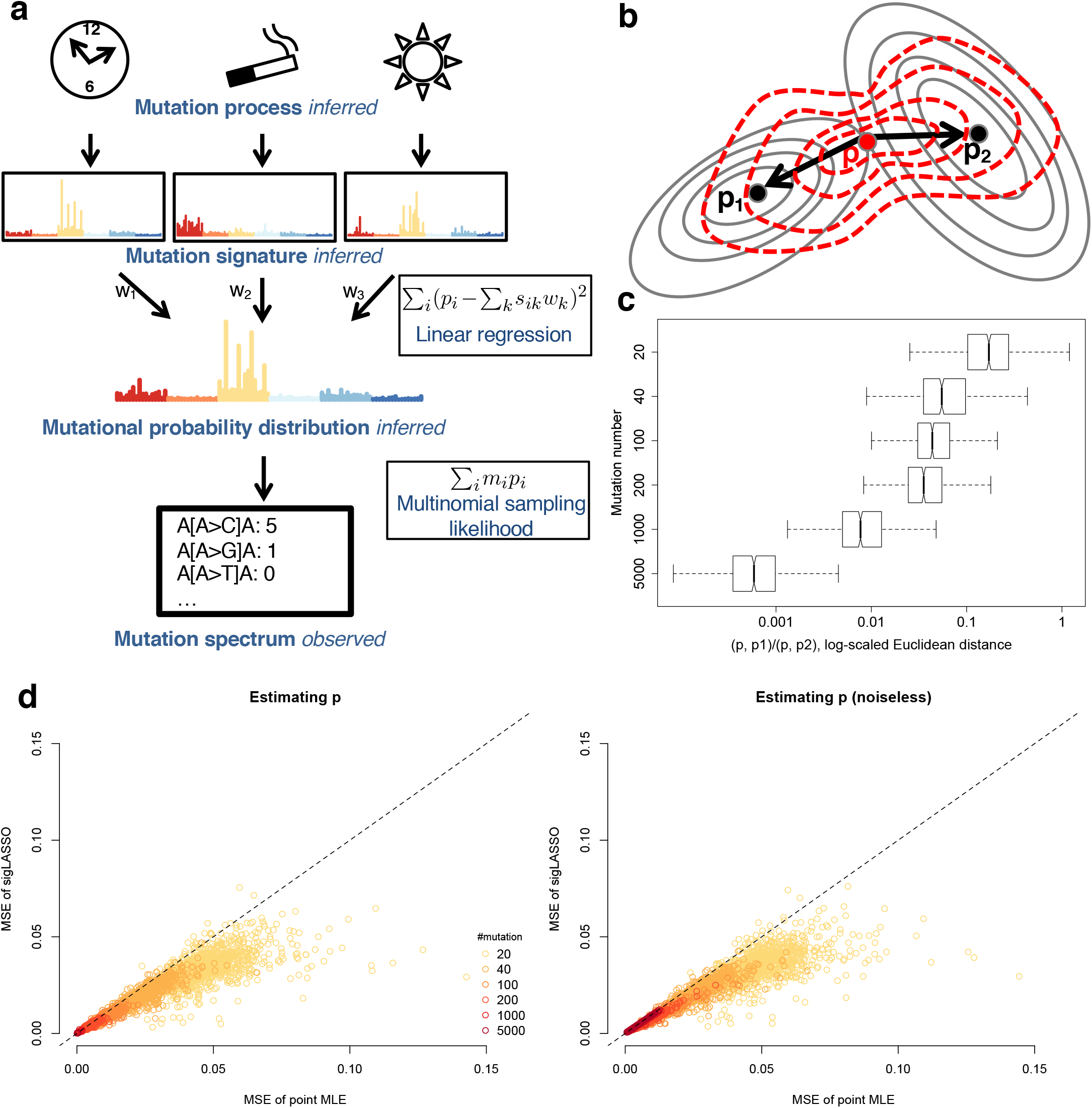
sigLASSO takes sampling variance into account. **(a)** A schematic graph showing the mixture model of mutational processes and signatures. **(b)** contour plot of the penalty function of multinomial sampling function (optimum at *p*_1_) and the least square of signature fitting (optimum at *p*_2_). sigLASSO tries to jointly optimize both penalties (red contour lines, optimum at *p*). **(c)** As mutation number increases, the inferred 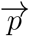 gets closer to the sampling MLE rather than the linear fitting as the sampling variance decreases. **(d)** The MSE of the estimation of 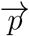 and the underlying noiseless signature mixture by sigLASSO and using the point MLE. Low mutation counts profiles benefit from sigLASSO the most. Priors were sampled uniformly from the ground true positives and negatives.

Then there are *K* latent mutational processes. Large-scale pan-cancer analyses identified *K* = 30 COSMIC signatures by NMF (with Frobenius norm penalty) and clustering.(2; 3) Here, our objective was to leverage the pan-cancer analysis and decompose mutations from new samples into a linear combination of signatures. Mathematically, we formulated the following non-negative regression problem, maintaining the original Frobenius norm:

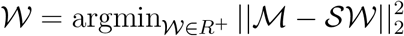

The mutation matrix, 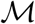 contains mutations of each sample cataloged into *n* trinucleotide contexts. 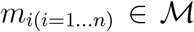 denotes the mutation count of the *i*^*th*^ category. 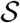 is a *n* × *K* signature matrix, containing the mutation probability in 96 trinucleotide contexts of the 30 signatures. 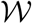 is the weights matrix, representing the contributions of 30 signatures in each sample.

### Sampling variance

In practice, this problem is optimized using continuous relaxation for efficiency and simplicity,(8) neglecting the discrete nature of mutation counts. This approach essentially transforms observed mutations into a multinomial probability distribution, making model estimation insensitive to the total mutation count. However, the total mutation count plays a critical role in inference. Assuming mutations are drawn from a latent probability distribution, which is the mixture of several mutational signatures, the mutations follow a multinomial distribution. The total mutation count is the sample size of the distribution, thus greatly affecting the variance of the inferred distribution.

For instance, 20 mutations within the 96 categories give us very little confidence in inferring the underlying mutation distribution. By contrast, if we observed 2,000 mutations we would have much higher confidence. Methods using continuous relaxation treat these two conditions indifferently. Here, we aimed to use a likelihood-based approach to acknowledge the sampling variance and design a tool sensitive to the total mutation count.

### sigLASSO model

We divided the data generation process into two parts. First, multiple mutational signatures mix together to form an underlying latent mutation distribution. Second, we observed a set of categorical data (mutations), which is a realization of the underlying mutation distribution. We used *m*_*i*_(*i* = 1 … *n*) to denote the mutation count of the *i*^*th*^ category. The vector 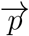 is the underlying latent mutation probability distribution with *p*_*j*_ denoting the probability of the *j*^*th*^ category. The total number of mutations is *N*.

To achieve variable selection and promote sparsity and interpretability of the solution, sigLASSO adds an L1 norm regularizer on the weights 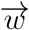 (i.e., coefficients) of the signatures with a hyper-parameter λ. LASSO is mathematically justified and can be computationally solved efficiently (14). Adding an L1 norm regularizer is equivalent to placing a Laplacian prior on 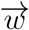 (19). 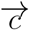 is a vector of *K* penalty weights (*c*_1_, *c*_2_, … *c*_*K*_), each indicating the strength to penalize the coefficient of a certain signature. This vector should be tuned to reflect the level of confidence in the prior knowledge. For example, a smaller penalty weight represents a stronger prior, reflecting higher confidence and vice versa.

Overall, from the generative model, we can determine the likelihood function for a single sample.

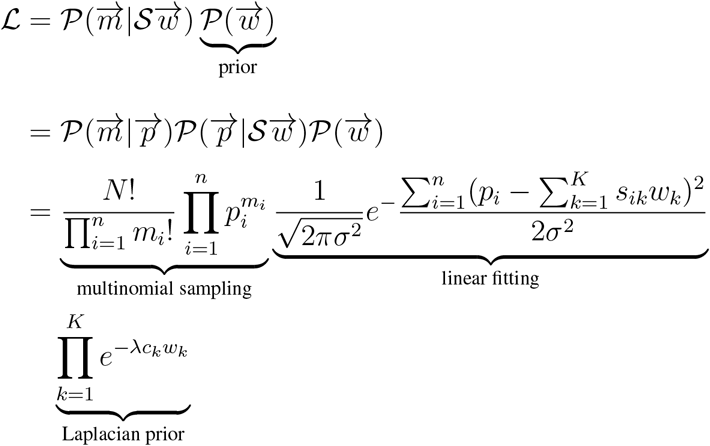

Our objective function is to maximize the log-likelihood function, which is given as:

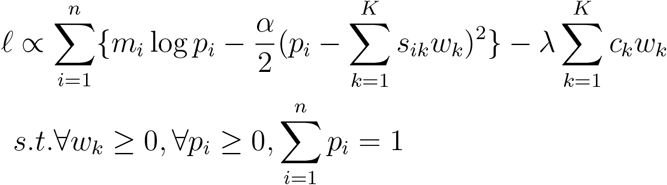

Here, *α* = 1/*σ*^2^. We can infer *α* from the residual errors from linear regression (see parameter tuning). Meanwhile, because of its continuous nature, 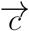 can be also effectively learned using patient information (e.g., smoking status, tumor size, or methylation status). We also used 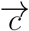 to perform adaptive LASSO (20) by initializing 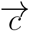 to 1/*β*_*OLS*_, where *β*_*OLS*_ are the coefficients from nonnegative ordinary least square. Our aim was to obtain a less biased estimator by applying smaller penalties on variables with larger coefficients.

### Optimizing sigLASSO

The negative log likelihood is convex in respect to both 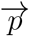 and 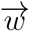 when evaluated individually. Hence, the loss function is biconvex. Instead of using a generic optimizer, we exploited the biconvex nature of this problem and effectively optimized the function by using alternative convex search, which iteratively updates these two variables (21).

**Algorithm 1.**
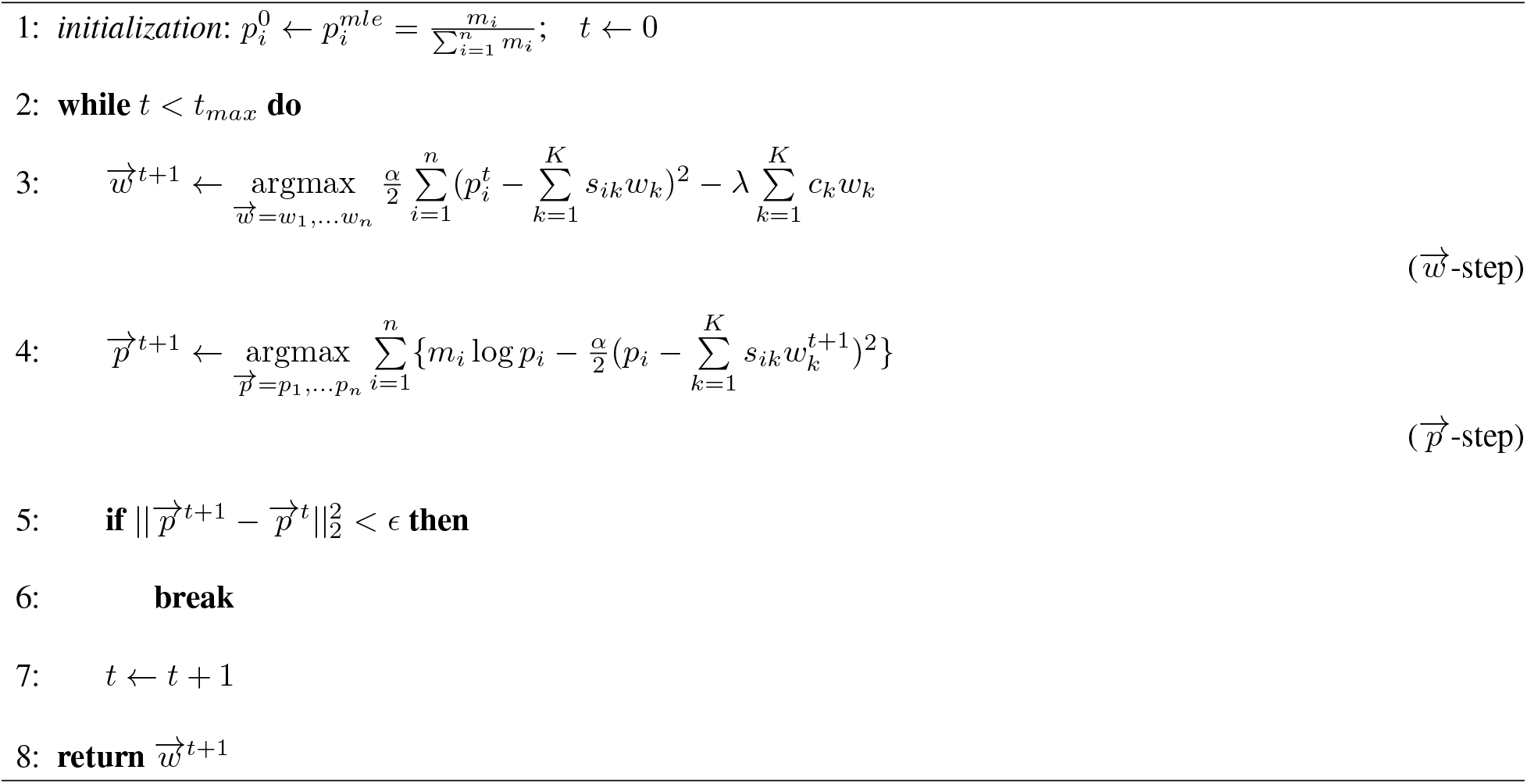
sigLASSO algorithm

Specifically, to begin the iteration, we initialized 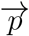 using maximum likelihood estimation (MLE). We started with the 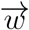–step, which is a nonnegative linear LASSO regression that can be efficiently solved by glmnet. (14) λ is parameterized empirically.

Next, we solved the 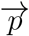 with a Lagrange multiplier to maintain the linear summation constraint 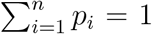. The nonnegative constraint of *p*_*i*_ is satisfied by only retaining the nonnegative root of the solution.

Intuitively, in the 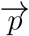–step, we tried to estimate 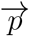 by optimizing the multinomial likelihood while constraining it to be not too far away from the fitted 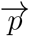. If we only used the point MLE of 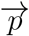 based on sampling and did not perform the 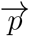–step, the model would assume the sampling is perfect and become insensitive to the total mutation counts. The trade-off in the 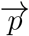–step between the multinomial likelihood and the square loss reflects the sampling error. The sampling size (sum of *m*_*i*_), the goodness of the signature fit (as reflected in *α*), and the overall shapes of 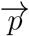 all affect the tension between sampling and linear fitting.

### Optimizing the 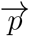 –step

In the 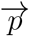 -step, we tried to solve the following problem with 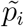 from the 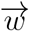–step.

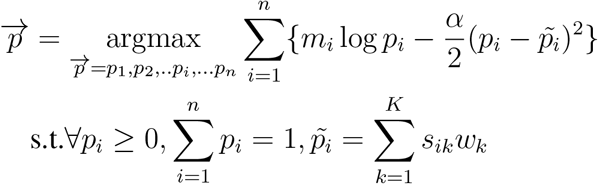

We added the Lagrangian multiplier Λ to satisfy the linear constraint of 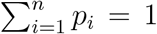 and took the derivatives with respect to *p*_*i*_(*i* = 1, 2…*n*) and Λ. This resulted in *n* + 1 equations.

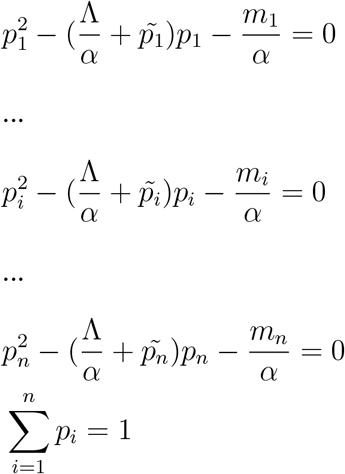

The roots of the first *n* quadratic equations are given by

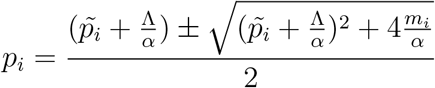

*α* = 1/*σ*^2^ is strictly positive and *m*_*i*_ is nonnegative. Therefore, if *m*_*i*_ = 0, there exists only one zero root and *p*_*i*_ = 0 iff. *m*_*i*_ = 0. If *m*_*i*_ > 0, there is exactly one negative and one positive root. Because we required ∀*p*_*i*_ ≥ 0, we only kept the positive root. The second derivative of the log-likelihood is 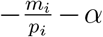, which is strictly negative. Therefore, the root we found is a nonnegative maximum.

We plugged all the roots into the last equation (i.e., the linear constraint) and used the R function uniroot() to solve Λ.

### Parameter tuning

We tuned λ by repeatedly splitting the nucleotide contexts into training and testing sets and testing the performance. Because mutations of the same single nucleotide substitution context are often correlated, we split the data set into eight subsets. Each subset contained two of each single nucleotide substitution. We then held one subset as the testing dataset and only fit the signatures on the remaining ones. After circling all eight subsets and repeating the process 20 times, we used the largest λ (which leads to a sparser solution) that gave an MSE 0.5 or 1 SD from the minimum MSE. λ was tuned whenever 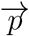 deviated far from the estimation from the previous round. By adaptively learning 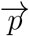, sigLASSO avoids overestimating the errors in the signature fitting and thus allows a higher fraction of mutations to be assigned with signatures. We fixed λ when the deviation was small to avoid the inherited randomness in subsetting affecting convergence. *α* = 1/*σ*^2^, *σ*^2^ is estimated using 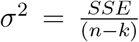, where k is the number of nonzero coefficients in the LASSO estimator and SSE is the sum of squared errors. (22) sigLASSO updates *α* after every LASSO linear fitting step. To avoid grossly overestimating *σ*^2^ (thus underestimating *α*) in the initial steps when 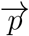 is far from the optimum, we set a minimum *α* value. In addition, because in practice factors such as unknown signatures violate the assumptions of linear signature regression, *α* tends to get overestimated. So we multiplied it by a confidence factor that represents how confident we are about the goodness of the signature fit. By default, we set min *α* = 400 and the confidence factor to 0.1. Users can further tune these values based on the strength of prior belief of noise level and confidence level of the signature model.

### Option for elastic net

Adding an L2 regularizer (i.e., elastic net) might improve and stabilize the performance when variables are highly correlated. (23) Therefore, we also implemented elastic net in sigLASSO. The objective function then became:

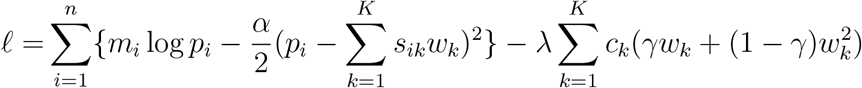

We tuned the hyperparameter *γ* by grid search together with λ. We always picked the largest *γ* and the largest λ (for better sparsity) that gave an MSE that was 0.5 SD from the minimal MSE. In simulations, we did not find that introducing L2 regularization added additional benefits. The model gave almost identical solutions to using L1 only. Nonetheless, we kept elastic net as an option in highly correlated signature scenarios to the user in our implementation.

### Data simulation and model evaluation

We downloaded 30 previously identified COSMIC signatures v2 (http://cancer.sanger.ac.uk/cosmic/signatures). We created a simulated dataset by randomly and uniformly drawing two to eight signatures and corresponding weights (minimum: 0.02). The reason for picking up to eight signatures is because 1) empirically, in cancer signature studies, most samples have only a few signatures. 2) We further confirmed the sparsity of signatures by running deconstructSigs on large-scale TCGA data and found 99.96% samples got assigned with eight or less signatures. 3) Moreover, biologically, we believe that it is unreasonable for more than 8 of the processes to act simultaneously in one sample. Many of these processes describe biological disjoint conditions, e.g. smoke, UV radiation and tissue-specific cellular processes.We simulated additive Gaussian noise at various levels with a positive normal distribution of up to 25 (1-25, uniformly drawn) randomly selected trinucleotide contexts. Then, we summed all the signatures and noise to form a mutation distribution. We sample mutations from this distribution with different mutation counts.

We ran deconstructSigs according to the original publication (8) and sigLASSO, both with and without prior knowledge of the underlying signature. To evaluate their performance, we compared the inferred signature distribution with the simulated distribution and calculated MSE. We also measured the number of false positive and false negative signatures in the solution (support recovery).

### Illustrating sigLASSO on 8,893 The Cancer Genome Atlas (TCGA) samples

To assess the performance of our method on real-world cancer datasets, we used somatic mutations from 33 cancer types from TCGA. We downloaded MAF files from the Genomic Data Commons Data Portal (https://portal.gdc.cancer.gov/). A detailed list of files used in this study can be found in supplemental table S1.

We extracted prior knowledge on active signatures in various cancer types from a previous pan-cancer signature analysis (http://cancer.sanger.ac.uk/cosmic/signatures).

### sigLASSO software suite

sigLASSO is implemented as an R package “siglasso”. It accepts processed mutational spectrums and VCF files. We provided functions to parse mutational spectrums from VCF files as well as some visualization methods in the package. “siglasso” allows users to specify biological priors (i.e., signatures that should be active or inactive) and their weights. “siglasso” uses 30 COSMIC signatures by default. Users are also given the option to supply customized signature files. It is computationally efficient; using default settings, the program can successfully decompose a whole genome sequencing (WGS) cancer sample in less than a few seconds on a regular laptop. For time profiling purposes, we ran siglasso and deconstructSigs on an Intel Xeon E5-2660 (2.60 GHz) CPU. We employed the R package “microbenchmark” to profile the function call siglasso() and whichSignatures(). For each setup, we generated 100 noiseless simulated data sets and repeated the process 100 times for each evaluation.

We made siglasso source code available at github.com/gersteinlab/siglasso.

### Evaluation criteria for signature assignment

One of the limitations of cancer signature research is that the ground truth of real samples typically cannot be obtained. Previous large-scale signature studies have relied largely on mutagen exposure association from patient records and biochemical knowledge on mutagenesis. Here, besides using simulation, we illustrated the outputs of different models and compared the results on real dataset with existing signature knowledge and distributions. Although no gold standard exists to evaluate the performance, we do have a few reasonable expectations about the solution:

#### Sparsity

One or more signature should be active in a given cancer sample and type. However, not all signatures should be active. A sparse distributed signature set among cancer samples and types is defined in previous *de novo* discovery studies. Any signature-fitting tool should follow and produce a similar signature distribution. Moreover, mutational processes are discrete in nature and tied with certain endogenous and environmental factors. An obvious example is that the UV signature should not exist in unexposed tissues. Existing signature-identifying methods aim to implicitly achieve sparse solutions by dropping signatures with small coefficients or pre-selecting a small signature subset for fitting.

#### The ability to abstain (decline to assign signatures to all mutations)

Signature assignment issues are often undefined due to collinearity of the signatures and a larger number of possible signatures. Overfitting is a serious concern, especially when the mutation counts are low (e.g., WXS or “silent” cancer genomes) or the fitting is poor (e.g., unknown mutational process or sample contamination causing high noise). A good solution should refuse to fully assign signatures to every mutation when it does not have enough confidence to do so and, instead, defer to the human researchers to decide.

#### Robustness

Solutions should be robust and reproducible. Signatures are not orthogonal, thus simple regression might lead to solutions that change erratically when a small perturbation is made in the observation. Moreover, the solution should reflect the level of ascertainment. Especially in WES, low mutation count is often a severe obstacle for assigning signatures due to undersampling. In particular, under low mutation count, not all of the operative signatures would be reliably discovered. It is better to abstain under high uncertainty.

#### Biological interpretability

The solution should be biological interpretable. Because of the biological nature of collinearity in the signatures, simple mathematical optimization might pick the wrong signature. Researchers now tackle this problem by simply removing the majority of predictors they believe to be inactive. sigLASSO allows users to supply domain knowledge to guide the variable selection in a soft thresholding manner, leaving space for noise and rare or unknown signatures. For instance, we expected to find divergent signature distributions in different cancer types, which is reported in previous *de novo* discovery studies. Various tissues have divergent endogenous biological features, are exposed to diverse mutagens, and undergo mutagenesis in dissimilar fashions. Signature patterns should be able to distinguish between cancer types. This can be achieved by using different cancer type specific priors.

These expectations are not quantitative, but they help direct us to recognize the most plausible solution as well as the less favorable ones.

## 3 RESULTS

### sigLASSO is aware of the sampling variance

By jointly optimizing both the sampling process and signature fitting, sigLASSO is aware of the sampling variance and infers an underlying mutational context distribution 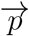. The underlying latent distribution is optimized in respect to both sampling likelihood and the linear fitting of signatures (Fig. 1b). In low mutation counts, the uncertainty in sampling increases and thus the estimated underlying distribution moves closer to the least square estimate (Fig. 1c). In contrast, when the total mutation count is high, the estimate of the distribution is closer to the MLE of the multinomial sampling process.

We illustrated how the mutation count affects the estimation of 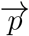 using a simulated dataset (five signatures, noise level: 0.1, see Methods). When the sample size was small (≤ 100), high uncertainty in sampling pushed the inferred underlying mutational distribution 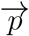 far from the MLE in exchange for better signature fitting. When the sample size increased, lower variance in sampling dragged 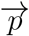 close to the sampling MLE and forced the signatures to fit even with larger error. Because linear fitting and sampling likelihood optimization mutually inform each other, concurrently learning an auxiliary sampling likelihood improves performance. We compared the accuracy of the estimation of 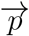 with and without this joint optimization (Fig. 1d). As expected, 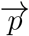 estimation in low mutation count performed worse. sigLASSO was able to achieve a lower MSE in estimating both 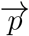 (with noise) and the underlying true signature mixture (noiseless).

### Performance on simulated datasets

We first evaluated sigLASSO on simulated datasets. Both sigLASSO (with and without priors) and deconstructSigs performed better with higher mutation number and lower noise (Fig. 2a, S1). A decrease in mutation number leads to an increase of uncertainty in sampling, which is mostly negligible in the high mutation scenarios. As expected, the MSE jumped to the 0.05-0.3 range regardless of the noise level when the mutation number was low. We observed a similar pattern in support recovery (i.e., precision/recall/accuracy). Thus, the error is dominated by undersampling rather than embedded noise. Despite giving a simpler, sparse solution, sigLASSO with priors outperformed sigLASSO with no priors and deconstructSigs in both MSE and support recovery. In support recovery, sigLASSO with priors showed a 5 − 10% gain in accuracy compared to deconstructSigs. sigLASSO with no priors also had better support recovery performance, measured by accuracy, than deconstructSigs. Overall, sigLASSO maintained a higher precision level when the mutation number decreased and/or the noise increased, which shows its ability to abstain (decline to assign signatures) and provide some control on the false positive rate. Notably, sigLASSO with priors maintained an accuracy above 0.8 in all simulation settings. Moreover, the precision of sigLASSO was minimally affected by the number of signatures.

**Figure 2:**
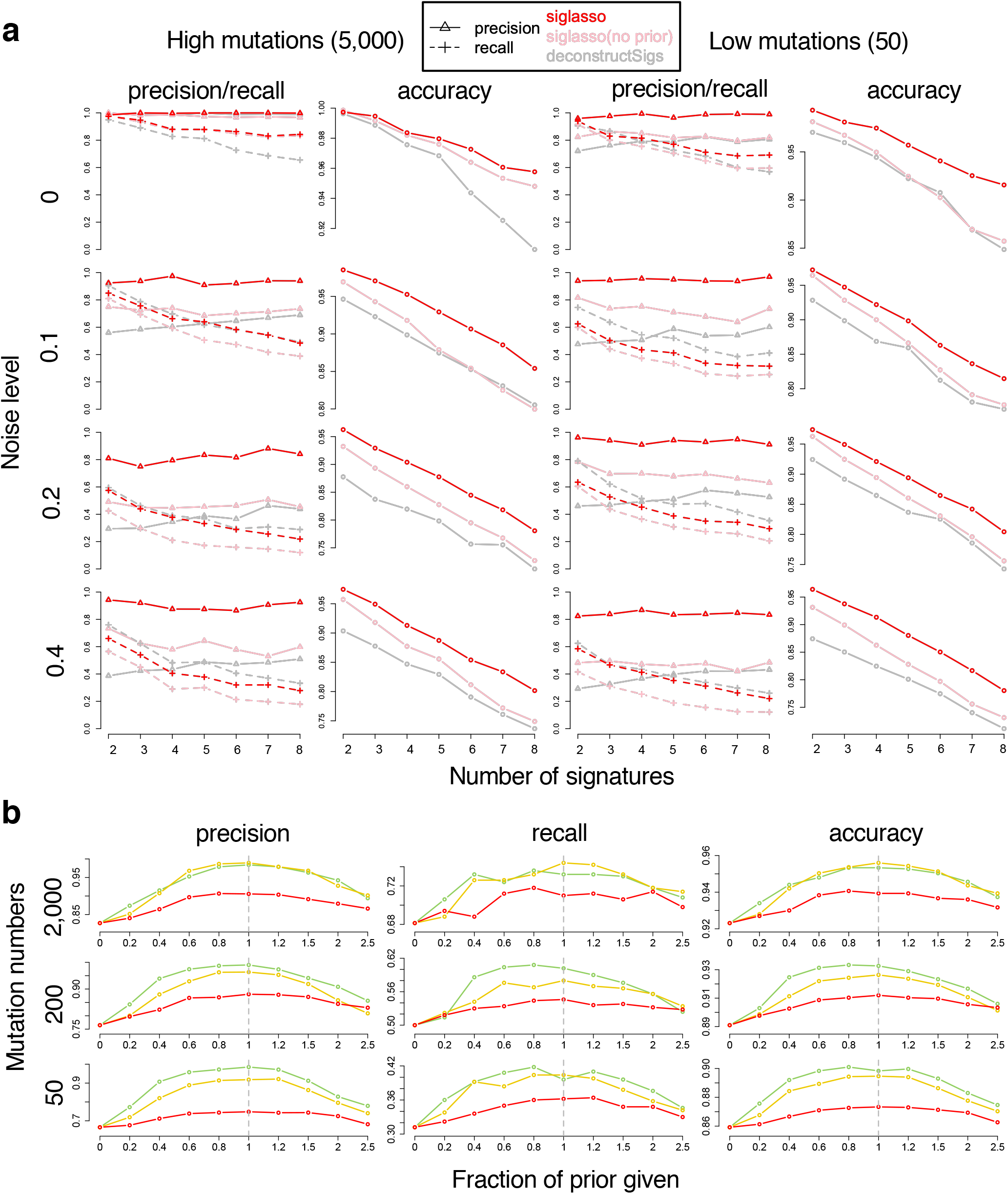
Performance on simulated datasets. **(a)** Support recovery on simulated datasets. Each setting is simulated 100 times. Penalty coefficients for priors: 0.1. **(b)** Support recovery on simulated datasets. Tuning the penalty weights using prior knowledge improves performance; x-axis: fraction of true signatures given as prior (larger than 1 indicates false signatures giving as priors). Penalty coefficients for priors: red, 0.5; yellow, 0.1; green, 0.01.

While the recall was slightly lower with sigLASSO than deconstructSigs in noisy settings, in very low and no noise situations sigLASSO achieved a higher recall due to its ability to adapt to different noise levels. By contrast, deconstructSigs, which assumes a fixed noise level, overly pruned the signatures in the post-fitting step when the noise was low. Lastly, adding correct priors helped boost both precision and recall significantly.

We next explored how different priors affect sigLASSO’s performance. Our experiments showed that using known signature as priors to tune the weights boosts performance. Priors improved performance even when we included only a small fraction of true signatures or blended in a large number of wrong signatures (Fig. 2b, S2). As the fraction of true signatures given as prior knowledge increased from zero, the performance immediately started improving and continued to do so. When more false signatures were mixed with true signatures given as prior knowledge, the performance slowly deteriorated. However, even with 1.5 times false signatures mixed in with true ones, the performance was slightly better (about 2% in accuracy) than the baseline that used no priors. Stronger priors had larger boosting effects to the solution, as expected.

### WGS scenario using renal cancer datasets

We next moved from synthetic datasets to real cancer mutational profiles, which are likely noisier than simulations and exhibit a highly non-random distribution of signatures.

We benchmarked the two methods using 35 WGS papillary kidney cancer samples (24). The median mutation count was 4,528 (range: 912-9,257). We found that without prior knowledge, both sigLASSO and deconstructSigs showed high contributions from signatures 3 and 5 (Fig. 3a, b). deconstructSigs also assigned a high proportion to signatures 8(9.9%), 9(6.9%), and 16(4.7%). Signatures 3, 8, 9, and 16 were not found to be active in pRCC in previous studies and currently no biological support connects them to pRCC (2). As expected, sigLASSO resulted in sparser solutions than deconstructSigs (mean signatures assigned: 3.40 and 4.43, respectively). Adding prior from COSMIC (Kidney cancer combined) helps sigLASSO to put more weight on Signature 5, and increase the number of signatures assigned to 4.06.

**Figure 3:**
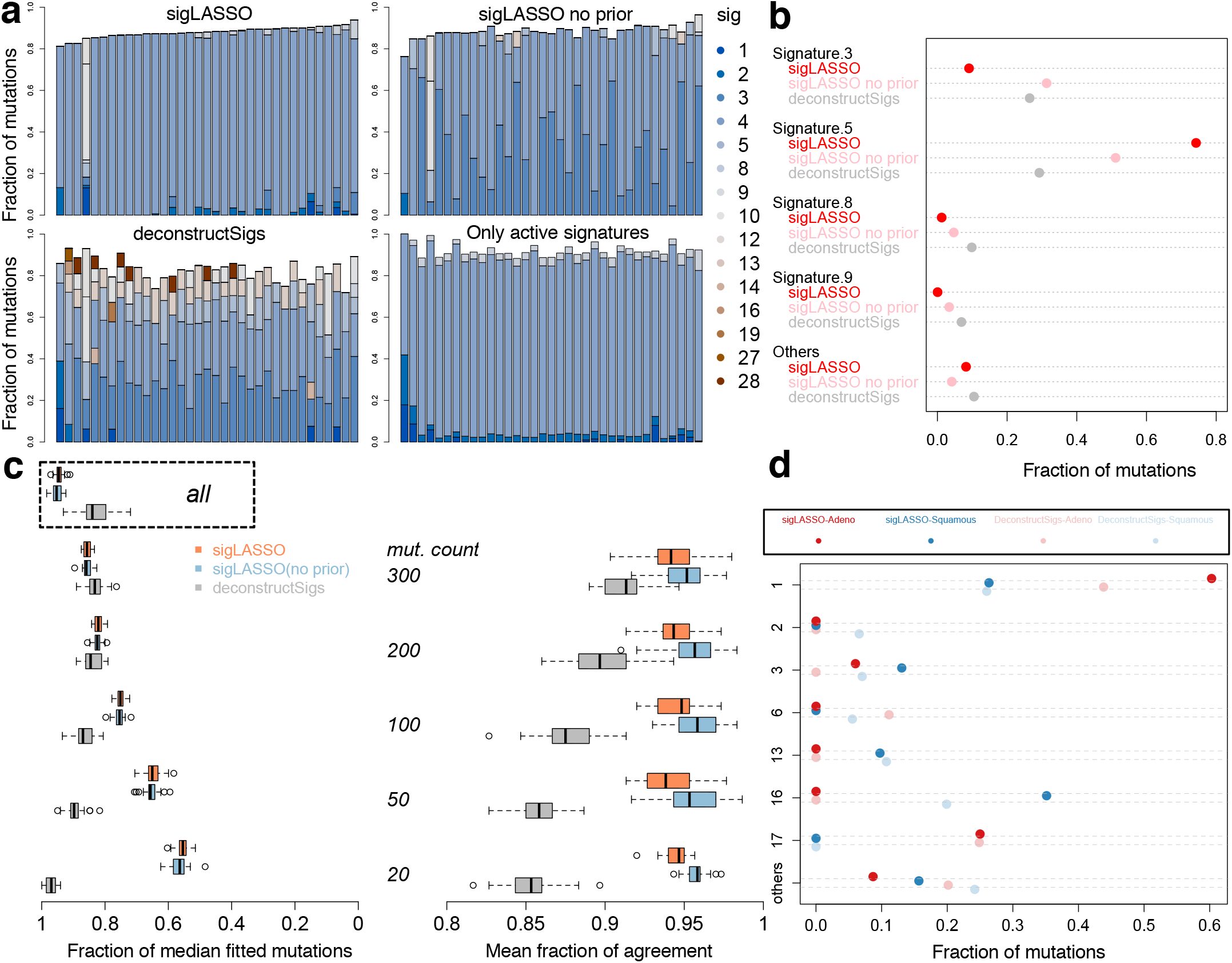
Performance on pRCC and ESCA samples. **(a)** Signature assignment for 35 WGS pRCC samples. Bar plots show the fractions of mutation signature assignment for each sample using sigLASSO, sigLASSO without prior knowledge and deconstructSigs, and simple nonnegative regression. **(b)** A dot chart showing the mean fraction of mutation signatures in each sample. Signatures that contributed less than 0.05 are grouped into “Others”. **(c)** Subsampling (10 times each size, with no replacement) of 30 WGS pRCC samples that have more than 3,000 mutations. Left panel shows the fraction of signature-fitted mutations. Results using all mutations are shown in box. Right panel shows the agreement (mean fraction of the consensus after binarizing whether a signature exists or not) among the methods. All *p* ≤ 2 × 10^−6^ (paired two-sided Wilcox test between sigLASSO (and sigLASSO, no prior) and deconstructSigs). **(d)** A dot chart showing the mean fraction of mutation signatures in each sample, grouped by two tools and histological subtypes (adenocarcinoma/squamous). Signatures that contributed individually less than 0.05 in all four cases are grouped into “others”.

However, if we naively subset the signatures and took the ones that were found to be active in previous studies, the signature profile was completely dominated by signature 5, to which only about 10% mutations on average were assigned with other signatures. Moreover, the model assigned highly similar signature profiles to all samples. This finding suggests an overly simple, underfitted model.

To show that sigLASSO is sensitive to fitting uncertainty, able to abstain, and is robust, we performed subsampling on 30 pRCC samples (mutation counts ≥ 3, 000, Fig. 3c). As the sample size decreased, sigLASSO assigned fewer mutations with signatures (mean fraction dropped from 0.93 (all mutations) to 0.55 (20 mutations)), reflecting the greater uncertainty in fitting. This quality allows the user to be aware of the uncertainty in the solutions.

Moreover, the model complexity also declined accordingly (Fig. S3). The mean number of signatures assigned by sigLASSO decreased from 4.06 to 1.67, representing simpler models. Last, the performance of sigLASSO was robust and stable, as evidenced by stable outputs even in low sampling counts. Multiple subsampling runs showed high agreement in active and inactive signatures (about 0.95 across all subsample sizes).

In contrast, deconstructSigs was unable to reflect model uncertainty. Surprisingly, as the mutation number decreased, the fraction of fitted mutations in deconstructSigs unexpectedly increased from 0.83 (all) to 0.97 (20 mutations). The model complexity, as reflected by the mean number of signatures assigned also increased from 4.43 to 4.70. The agreement between subsamples dropped significantly as the subsampling size decreased, indicating the solution is unstable and potentially overfitted.

### WES scenario using esophageal carcinoma datasets

We next aimed to evaluate the two methods on 182 WES esophageal carcinoma (ESCA) samples with more than 20 mutations. The median mutation count was 175.5 (range: 28-2,146), which is considerably lower than WGS but typical for WES.

In sigLASSO, the L1 penalty strength was tuned based on model performance. In a low mutation count setting, ESCA WES dataset, the model variance was high, which pushed up the penalty. As expected by its ability to abstain, sigLASSO assigned a lower fraction of mutations with signatures than deconstructSigs (median: 0.68 and 0.90; interquartile range: 0.56-0.75 and 0.86-0.94, respectively, Fig. S4). deconstructSigs did not achieve 100% assignment because it performed a hard normalization of the coefficients after an unconstrained binary search and then discarded signatures that had ≤ 6% contribution. The leading signatures were 1, 16, 17, 3 and 13 in sigLASSO and 1, 17, 16, 6, 13, 3 and 2 in deconstructSigs. deconstructSigs has been applied to distinguish between two different histological types of esophageal cancer (8). We demonstrated that sigLASSO generates a sparser but comparable result with wider signature 1, and 16 gaps between the subtypes (Fig. 3d, S5). The adenocarcinoma subtypes had higher fractions of signature 1 and 17, and lower fractions of signature 3, 13 and 16.

### Performance on 8,893 TCGA samples

We ran sigLASSO and deconstructSigs with step-by-step set-ups on 8,893 TCGA tumors (33 cancer types, Table S11) with more than 20 mutations (Fig. 4, S6). The median mutation number is 100, interquartile range (IQR) is 52-200.

**Figure 4:**
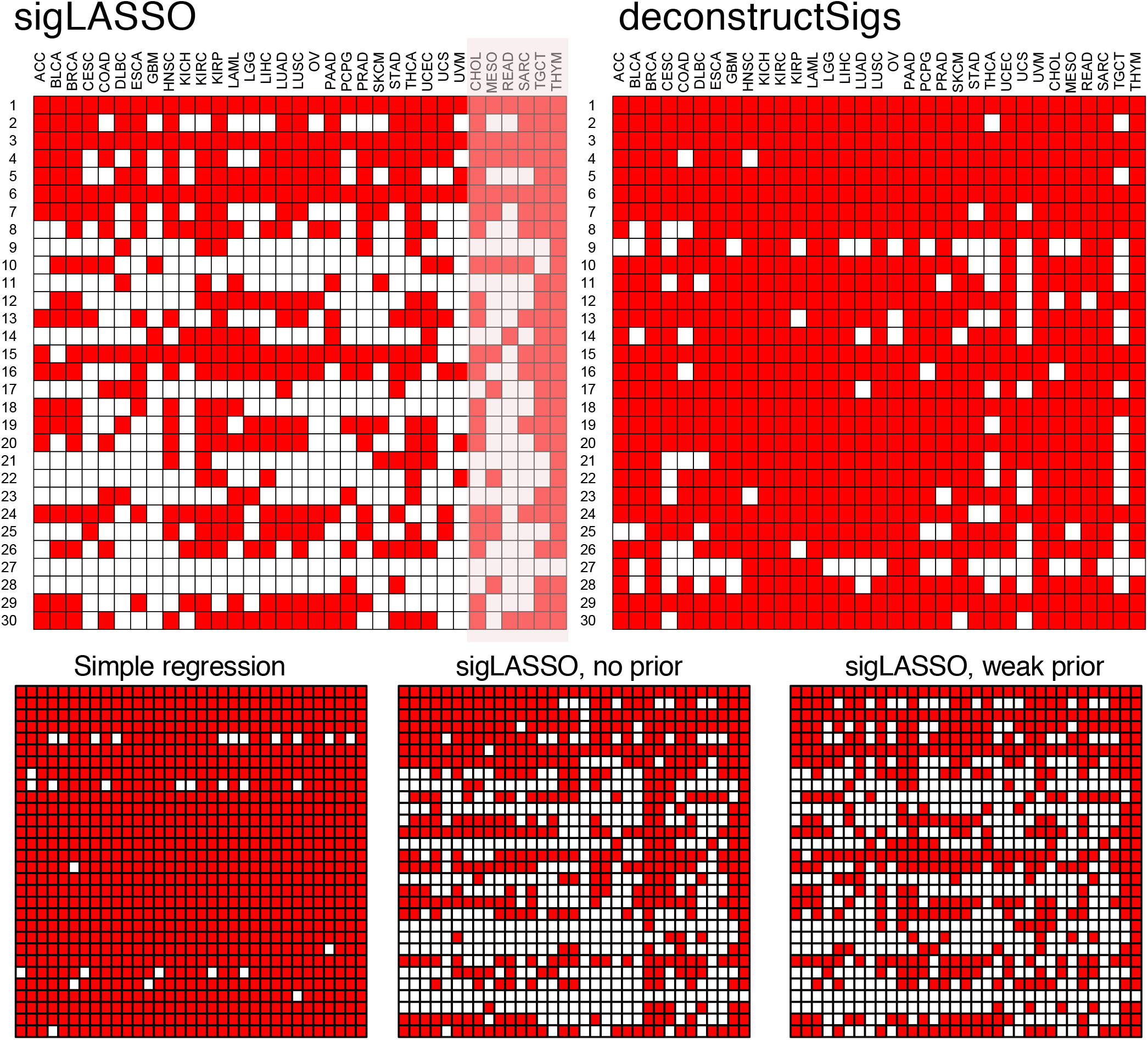
Performance on 33 TCGA cancers. Active signatures (total contribution > 0.1%) in 33 cancer types using different methods. Only 27 cancer types have previously known signature distributions (others are shaded). Penalty coefficients of priors used for sigLASSO: 0.01. Weak priors: 0.5. Priors were taken from COSMIC, and provided with our “sigLASSO” R package.

Simple nonnegative regression resulted in an overly dense matrix. Applying an L1 penalty made the solution remarkably sparse. Then, by incorporating the prior knowledge, the signature landscape further changed without significantly affecting the assignment sparsity. The change was inconsistent with the priors given. With low-count WES data (upper quartile: 200 and 96 mutational contexts), we expect the signature assignment to be very sparse as the uncertainty is high, the model should from making assignments. sigLASSO indeed assigned low number of signatures to the samples (median number of signatures assigned per sample: 1, IQR: 1-2). In comparison, the solutions of deconstructSigs were less sparse (median number of signatures assigned per sample: 4, IQR: 3-5). 18.5% samples were fitted by deconstuctSigs with six signatures or more, compared to 0.4% by sigLASSO. We provide a kidney cancer example and detailed paired analysis in the Supplement (Fig. S7, 8).

When the mutation number cut-off increases to 50, the above results of the two methods still stand (Fig. S9). We also released all of the signature results of 8,893 TCGA samples on the sigLASSO GitHub site.

Using the large-scale tumor signature profiles, we further explored the correlation of smoking signature and smoking status using annotations from a previous study (15). In lung adenocarcinoma (LUAD) samples, smoking samples carry significantly higher signature 4 (“smoking signatures”) fractions than non-smoking ones (median: 0 and 0.68, respectively, *p* ≤ 1 × 10^−15^, Wilcoxon rank test, Fig. S10). Similarly, in lung squamous cell carcinoma (LUSC) samples, we observed high fractions of signature 4 in smokers, but because only 3.5% of the LUSC cohort are nonsmokers (6/171), we were underpowered to draw a statistical significance. In non-lung cancer samples (N = 1,500), we also found a weaker but statistically significant trend of higher signature 4 in smokers (mean: 0.008 and 0.038, respectively, *p* = 3.3 × 10^−8^, Wilcoxon rank test, Fig. S10). This result is in agreement with previous studies on smoking signatures (15)

### sigLASSO is computationally efficient

sigLASSO iteratively solves two convex problems. The 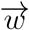−step can be solved using a very efficient coordinate descent algorithm (glmnet) (14). The 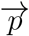−step is solved by a set of quadric equations. We observed empirically that the solution quickly converges in a few iterations even with low mutation numbers. Meanwhile, deconstructSigs uses binary search instead of regression to try every coefficient by looping through all signatures at each iteration.

By profiling sigLASSO and deconstructSigs (Fig. 5), we noticed that neither total mutation numbers nor signature numbers remarkably affected the running time of sigLASSO. With a high mutation number, sigLASSO was 3-4 times faster than deconstructSigs; with a low mutation number (50 mutations), these two tools showed a comparable computation time. Noticeably, despite using less time sigLASSO employs empirical parameterization and alternative optimization, which regresses the signature fitting problems hundreds of thousands of times with different parameters in a typical run. Therefore, by carefully designing an effective algorithm, the core fitting step of sigLASSO is orders of magnitudes faster than deconstructSigs. This enables sigLASSO to probe the data complexity and accordingly tune the model complexity.

**Figure 5:**
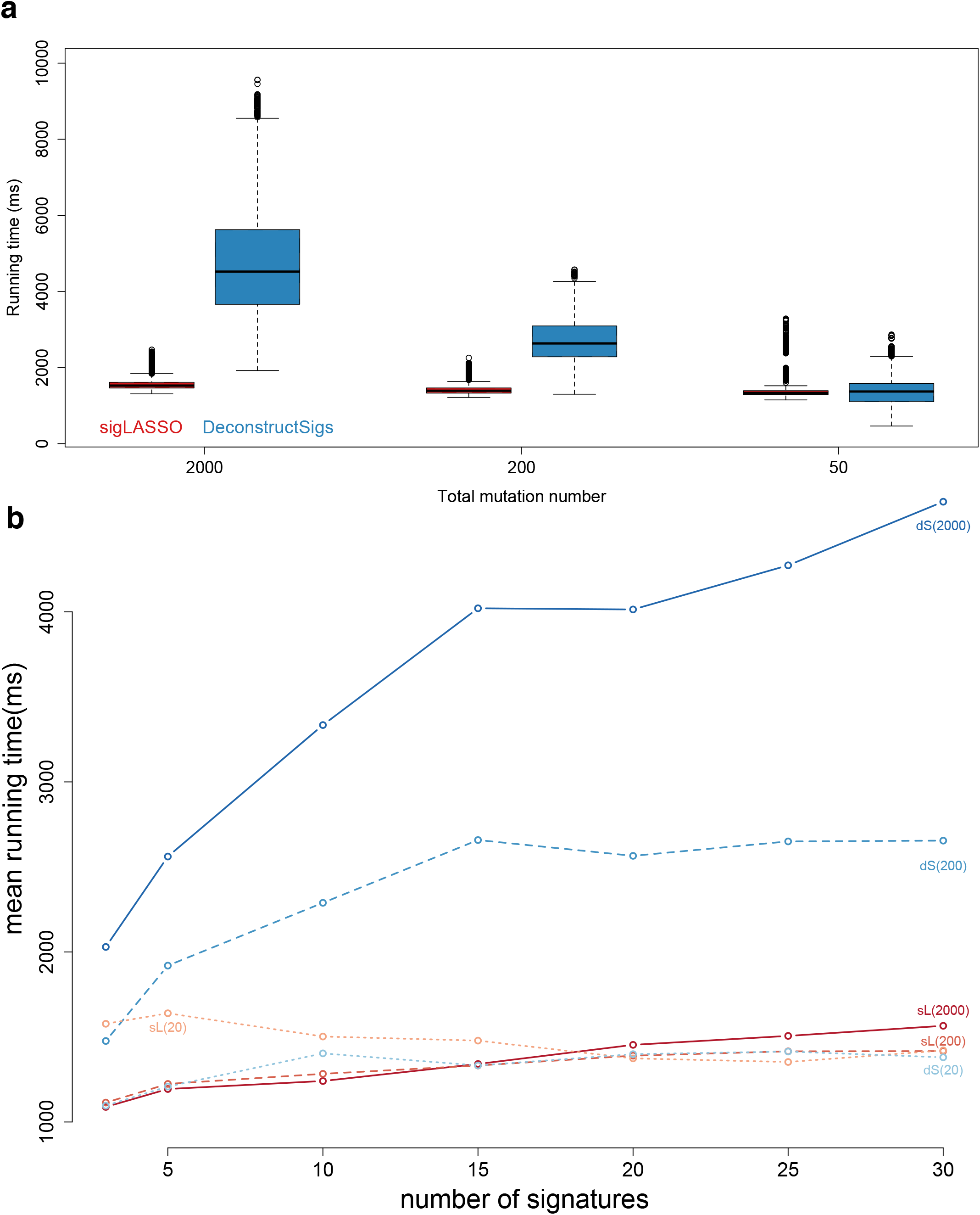
**(a)** Running time of sigLASSO and deconstructSigs at different total mutations numbers **(b)** Running time of sigLASSO at different numbers of signatures (downsampled from 30 COSMIC signatures) simulated with three different mutations numbers (in parenthesis). Noise level: 0.1 sL: sigLASSO, dS: deconstructSigs

## 4 DISCUSSION

Studies decomposing cancer mutations into a linear combination of signatures have provided invaluable insights into cancer biology.(4; 5; 6; 25) Indeed, researchers have gained a better understanding of one of the fundamental driving forces of cancer initiation and development, mutagenesis, by inferring mutational signatures and latent mutational processes.

A practical problem for many researchers is how to leverage results from large-scale signature studies and apply them to a small set of incoming samples. Although this might seem to be a simple linear system problem, core challenges include (1) preventing over- and underfitting on only one sample, often with very few mutations (especially in WES), (2) achieving proper variable selection from a large amount of recognized signatures, and (3) promoting interpretability.

First, with less than a few hundred mutations, sampling variance becomes a significant factor in reliably identifying signatures. Therefore, the fitting scheme should be aware of the sampling variance, which is especially pronounced in low mutation count scenarios (WES or cancer types with low mutation burden). Ideally, the tool should be able to attribute the signatures by flexibly inferring the underlying true mutation distribution given the sampling variance and the signature fitting performance. Second, the large number of available signatures necessitates proper variable selection, especially with a limited amount of observed data. Signature studies on large-scale cancer datasets have revealed that mutational signatures are not all active in one sample or cancer type; in most tumor cases, only a few signatures prevail. A recent signature summary suggested that 2 to 13 known signatures are observed in a given cancer type (based on 30 COSMIC signatures), which might include hundreds and even thousands of samples. Therefore, sparse solutions are in line with previous *de novo* studies, and are biologically sound and interpretable. In addition, sparse solutions can give better predictions due to lower estimator variance. Third, a desirable method should be parameterized according to the data complexity to achieve optimum fitting. Finally, mutational signatures are not orthogonal due to their biological nature. Collinearity of the signatures will lead to unstable fittings that change erratically with even a slight perturbation of the observation. deconstructSigs was the first tool that could identify signatures even in a single tumor. Instead of regression, this tool uses binary search to iteratively tune coefficients. To achieve sparsity, deconstructSigs performs post-hoc pruning with a preset 6% cut-off value. The mutation spectrum is normalized before fitting, thus making mutation counts invariant to the model. Moreover, the binary search operates on unbounded coefficients and uses a hardcoded upper and lower bounds, which provides no guarantee of finding the optimal solution. Finally, the greedy nature of stepwise coefficient tuning is prone to eliminating valuable predictors in later steps that are correlated with previously selected ones.(26)

Here, we describe sigLASSO, which simultaneously optimizes both the sampling process and an L1 regularized signature fitting. By explicitly formulating a multinomial sampling likelihood into the optimization, we designed sigLASSO to take into account the sampling variance. Meanwhile, sigLASSO uses the L1 norm to penalize the coefficients, thus achieving effective variable selection and promoting sparsity. By fine-tuning the penalizing terms using prior biological knowledge, sigLASSO is able to further exploit previous signature studies from large cohorts and promote signatures that are believed to be active in a soft thresholding manner.

Jointly optimizing a mutation sampling process enables sigLASSO to be aware of the sampling variance. By additionally modeling an auxiliary multinomial sampling process and corresponding distribution, we demonstrated that sigLASSO achieves better and more stable signature attribution, especially in cases with low mutation counts. In cancer research, WES data is abundant, but it also suffers from undersampling in signature attribution. In these cases, sigLASSO generates more reliable and robust solutions. We showcase the successful application of sigLASSO on 8,893 TCGA WES samples. Overall, sigLASSO achieves better sparsity than deconstructSigs and promote structures in the previous signature studies by injecting priors into the optimization process. The final signature distribution of neither sigLASSO nor deconstructSigs is identical to the original signature studies, which is probably due to biases in the original signature discovery, sequencing/variants calling, and unknown hypermutational processes. We noticed deconstructSigs assigned signatures to 100% of the mutations in most samples, while in sigLASSO assignment fraction correlated with cancer types. This abstain pattern indicates the presence of potentially unknown signatures and cancer subtypes.

As the cost of WGS drops rapidly, we expect an even greater number of cancer samples to be sequenced.(27) The vast amount of cancer genomics data will give scientists larger power to discern unknown or rare signatures. The growing number of signatures will eventually make the signature matrix underdetermined (when *k* > 96, i.e., the number of possible mutational trinucleotide contexts). A traditional simple solver method would give infinitude (noiseless) or unstable (noisy) solutions in this underdetermined linear system. However, by assuming the solution is sparse, we were able to apply regularization to achieve a simpler, sparser, and more stable solution. Furthermore, sigLASSO is very efficient and thus able to handle a larger number of samples and signatures, as well as hyperparameter optimization.

Moreover, sigLASSO does not specify a noise level explicitly beforehand, but instead empirically tunes parameters based on model performance. By contrast, deconstructSigs specifies a noise level of 0.05 to derive a cut-off of 0.06 to achieve sparsity. In general, sigLASSO lets the data itself control the model complexity and leave any post-hoc filtering to users. Abstention is a desired trait in machine learning models. In addition to the objectives, the model learns what it knows and what it does not (“KWIK: Knows What It Knows”),(28) In both subsampling and simulation experiments, sigLASSO is able to abstain under high uncertainty, providing solutions with consistently high precision and robustness under various scenarios.

Next, due to the collinearity nature of signatures, pure mathematical optimization might lead algorithms to select wrong signatures that are highly correlated with truly active ones. To overcome this problem, sigLASSO allows researchers to incorporate domain knowledge to guide signature identification. This input could be cancer-type-specific signatures or patient clinical information (e.g., smoking history or chemotherapy). Furthermore, we transformed the binary classification (“active” or “inactive”) to a continuous space, with weights indicating the strength of the prior. These weights could be effectively learned using patient information or other assays (e.g., RNA sequencing or methylation arrays). Moreover, sigLASSO can adapt to sparsity promoting schemes in *de novo* discovery tools through priors. For example, SparseSignatures((12)) also uses L1 regularization but does not penalize on “background signatures”. sigLASSO could use this piece of information in its priors to better assign signatures when working together with SparseSignatures. We showcased the performance of sigLASSO on real cancer datasets. Although we lack the ground truth of the operative mutational signatures in tumors, we have several reasonable beliefs about the signature solution. sigLASSO produced signature solutions that are biologically interpretable, properly align with our current knowledge about mutational signatures, and well distinguish cancer types and histological subtypes.

We also implemented elastic net with hyperparameter optimization in sigLASSO. Elastic net blends L2 with L1 regularization and in principle demonstrates better performance and stability than LASSO on strongly correlated features.(23) We found that elastic net did not improve the performance (measured as meaningful reduction in MSE in cross-validation) in our simulations using the current 30 COSMIC signatures, likely because the correlations between them are not too high. However, with more cancer samples sequenced every day, researchers will gain power to discern highly correlated mutational processes and grow the size of the signature set significantly. Therefore, elastic net might be beneficial in the near future.

Finally, sigLASSO uses quadratic loss, which follows the previous *de novo* studies.(2; 3; 5) By doing so, sigLASSO is unbiased in detecting signatures in cancer samples. Nonetheless, the initial discovery suffers from several limitations. First, the actual number of mutational processes is likely higher than 30, the current number of COSMIC signatures. This is because the power of the *de novo* discovery is bounded by the amount of available data. Second, the nature of mutagenesis leads to some mutational processes being more prevalent than others. Some of the signatures are cancer type specific. The original signature discovery did not use a balanced dataset with equal numbers of cancer types. Third, the original NMF objective function employs quadratic loss, which might lead to bias towards “flat” signatures over “spiky” ones. In addition to the original NMF approach, researchers have proposed other decomposition methods for signature discovery. For example, SomaticSignatures(29) uses PCA. Because the loss function is also quadratic, we expect sigLASSO to work seamlessly with SomaticSignatures.

Our simulation reveals that sigLASSO does not have a discernable preference to assign to either spiky or smooth signatures (Fig. S12). However, we found that the signature distribution from sigLASSO and deconstructSigs cannot be fully explained by the current signature knowledge. Further research probing the biological foundation of the signatures and quantifying the biological priors will help to advance our understanding and improve the interpretability of signature assignments.

sigLASSO exploits the previously overlooked mutation sampling uncertainty and formulates a framework that jointly optimizes the objective (“signature fitting with L1 regularization”) and the sampling likelihood. In biological experiments, low-count observations on discrete variables are common. For example, in single cell RNA sequencing (scRNAseq), the measured discrete mRNA counts are often very low or even zero (due to under-sampling). Our joint optimization approach could have further implications in these scenarios. We, indeed, find some similar work in scRNAseq.(30; 31) Another method for *de novo* signature discover, EMu, formulates the discreet mutational process as a Poisson generative model. (32) When the total mutational count is fixed, such Poisson generative model is equivalent to our multinomial sampling process. Our work also could be extended to mutagenesis modeling and parameter estimation—for example, in estimating the nucleotide-specific background mutation rate in cancer.

## 5 ACKNOWLEDGEMENTS

This work was supported in part by AL Williams Professorship and the facilities and staffs of the Yale University Faculty of Arts and Sciences High Performance Computing Center. We thank Harrison Zhou, Cong Liang, Jinrui Xu, Diego Galeano, and Mengting Gu for helpful discussion.

